# Integrating AlphaFold and deep learning for atomistic interpretation of cryo-EM maps

**DOI:** 10.1101/2023.02.02.526877

**Authors:** Xin Dai, Longlong Wu, Shinjae Yoo, Qun Liu

## Abstract

**Interpretation of cryo-electron microscopy (cryo-EM) maps requires building and fitting 3-D atomic models of biological molecules. AlphaFold-predicted models generate initial 3-D coordinates; however, model inaccuracy and conformational heterogeneity often necessitate labor-intensive manual model building and fitting into cryo-EM maps. In this work, we designed a protein modelbuilding workflow, which combines a deep-learning cryo-EM map feature enhancement tool, CryoFEM (Cryo-EM Feature Enhancement Model) and AlphaFold. A benchmark test using 36 cryo-EM maps shows that CryoFEM achieves state-of-the-art performance in optimizing the Fourier Shell Correlations between the maps and the ground truth models. Furthermore, in a subset of 17 datasets where the initial AlphaFold predictions are less accurate, the workflow significantly improves their model accuracy. Our work demonstrates that the integration of modern deep learning image enhancement and AlphaFold may lead to automated model building and fitting for the atomistic interpretation of cryo-EM maps.**

## Introduction

The resolution revolution in cryo-electron microscopy (cryo-EM) has greatly accelerated the experimental determination of protein structures (*1*). The reconstructed cryo-EM maps have reached near-atomic resolutions. However, to make use of cryo-EM maps to understand protein function, atomistic protein structures, including chain tracing, amino-acid residue registration, side-chain conformation, and 3-D coordinates, have to be modeled and refined.

The construction of atomic models from cryo-EM maps is a complex undertaking. When the resolution of these maps reaches 3.5 Å resolution or better, software tools such as PHENIX (*2*) or BUCCANEER (*3*) are typically employed to construct partial models. Subsequent model refinement and validation necessitate iterative adjustments, a time-consuming process typically carried out using programs like Coot (*4*) and PHENIX.Refine (*5*). Within this modeling process, the identification of secondary structure features like *α*-helices and *β*-strands in a cryoEM map forms a crucial initial step. To aid in this secondary structure recognition, machine learning methods have been integrated into software tools like SSELearner (*6*), *γ*-TEMPy (*7*), EMNUSS (*8*), and Emap2sec (*9*), automating the detection of these key structural elements. Deep learning methodologies, especially those utilizing 3D convolutional neural networks such as UNet (*10*), have made significant strides in recent years, enabling the prediction of protein backbone structure. C-CNN (*11*) and SEGEM (*12*) serve as prominent instances of this deeplearning-based approach. Further advancements in this direction have led to the development of sophisticated tools including DeepMM (*13*), DeepTracer (*14*), CR-I-TASSER (*15*), and ModelAngelo (*16*).

A protein’s sequence determines its structure and function. Recent advances in deep learning algorithms have made a breakthrough in protein structure prediction with representatives of AlphaFold (*17*), RoseTTAFold (*18*), ESMFold (*19*) and OmegaFold (*20*). These programs predict high-quality atomic models that can be used to support experimental protein structure determination (*21*). Among these programs, AlphaFold produces the most accurate atomic models (*19, 20*). AlphaFold takes the sequence of a protein to predict a 3-D atomic model that can serve as a starting point for the interpretation of cryo-EM maps through model building and refinement. However, the AlphaFold model represents a still structure with knowledge learned from the experimentally determined structures deposited to the Protein Data Bank (PDB) (*22*). The AlphaFold-predicted models may have poor quality and do not accommodate conformational changes captured in cryo-EM. It is thus highly desirable to have a model-building workflow that leverages AlphaFold to facilitate atomistic interpretations of cryo-EM maps.

AlphaFold can use structure templates to improve its prediction accuracy. Recently, Terwilliger *et. al* (*23*) proposed an intuitive solution, which first fits and refines the AlphaFold model against its cryo-EM map, then uses the refined model as a template to guide AlphaFold prediction. They demonstrate that refined templates help improve the quality of the predicted models. However, most of the proteins tested in Ref. (*23*) are relatively small, averaging 311 residues. For several examples, their strategy resulted in model overfitting.

Cryo-EM maps are averaged 3-D reconstructions, each from thousands to millions of particles. Due to radiation damage, heterogeneity, and intrinsic molecular motion, the high-resolution contrast in cryo-EM densities is lost, causing blurring. Therefore, a contrast-restoration process using a global B-factor sharpening has been routinely used to improve map interpretability (*2, 24, 25*). However, a global B factor can not accommodate local radiation damage, protein heterogeneity, and flexibility. Consequently, local B-factor sharpening methods were developed to post-process cryo-EM maps as implemented in programs LocScale (*26*), localDeblur (*27*), and Locspiral (*28*). By learning structural features from atomic coordinates, deep learning methods were also deployed to perform local B-factor sharpening for cryo-EM (*29–32*). Among these deep-learning methods, DeepEMhancer (*32*), which is based on a 3D UNet model (*10*) to improve the cryo-EM density map features is one of the popular tools to enhance cryo-EM maps.

In this study, we designed a deep learning model, CryoFEM (**Cryo**-EM **F**eature **E**nhancement **M**odel), which can effectively enhance the features in the cryo-EM maps. We developed a workflow to integrate the CryoFEM and AlphaFold to facilitate the model-building and refinement processes. We show that our CryoFEM outperforms DeepEMhancer by generating less noisy density maps that have higher correlations with the associated protein models. In addition, our model-building workflow allows the refinement of larger proteins (chain length of 1017 residues on average) and significantly improves the atomistic interpretability of cryo-EM maps where their initial AlphaFold predictions are less accurate.

## Results

### Map to model workflow

Our protein model-building workflow is illustrated in Fig. 1. The input is the target sequence and its associated raw cryo-EM map. The features in a raw map is enhanced by CryoFEM with its model structure shown in Fig. S1 and Table S1. The output is the atomic coordinates that have been modeled and fitted into the CryoFEM-enhanced map. Our model-building strategy follows that of Ref. (*23*), but we used the CryoFEM-enhanced map for rebuilding AlphaFold models. The workflow can be iterated multiple times, although the gain will likely be marginal after the first cycle (*23*).

**Figure 1:**
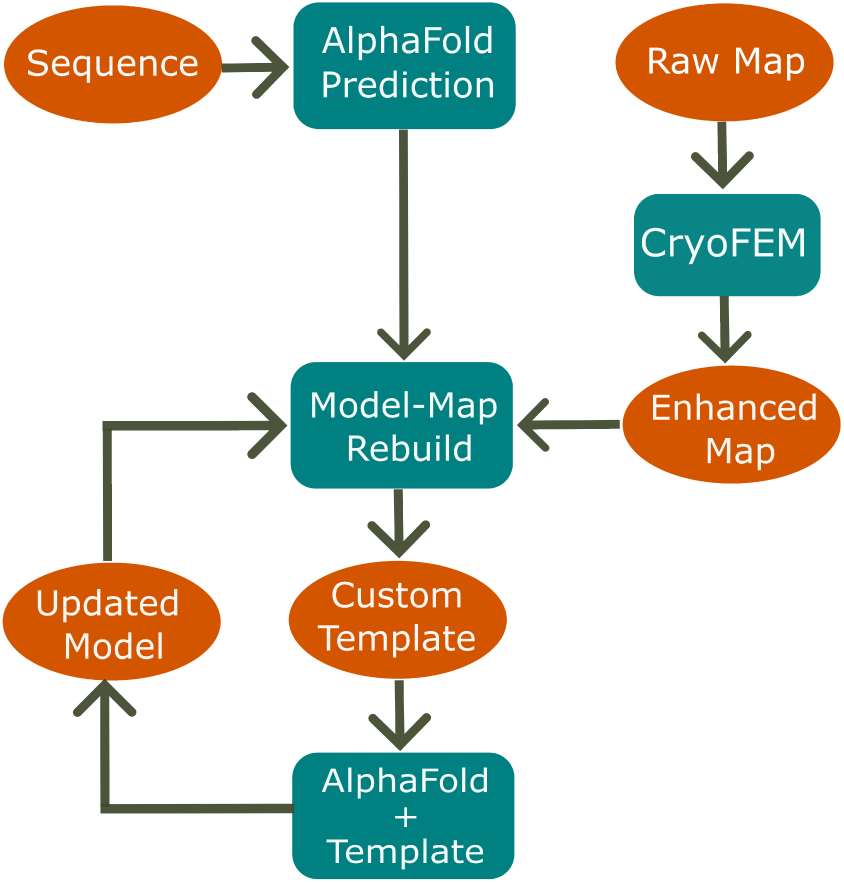
Schematics of the map to model workflow. An initial model is from AlphaFold and the enhanced cryo-EM map is from our deep-learning model CryoFEM. *phenix.dock and build* is to rebuild the model with the CryoFEM-enhanced density map. The resulting model serves as a template for the next round of AlphaFold prediction and *phenix.dock and build* model building, completing a full cycle of the workflow.

### Cryo-EM map enhancement

Upon training the CryoFEM model to convergence (Fig. S2), we compared the performance of CryoFEM with DeepEMhancer and two commonly used non-deep learning map sharpening methods, namely, *phenix.auto sharpen* (*33*) (global B-factor sharpening) and the local B-factor sharpening tool LocScale (*26*) using 36 cryo-EM maps, including 20 maps previously utilized by DeepEMhancer. The reported resolutions of these maps span from 2.8 to 6.7 Å, with a median resolution of 3.45 Å. Table 1 presents the median values for four distinct real-space model-map correlation coefficients calculated using *phenix.map model cc*. Our method, CryoFEM, noticeably surpasses the raw maps in terms of CC_box_ and CC_peak_ values, while demonstrating comparable values for CC_volume_ and CC_mask_. Table 2 enumerates the median resolutions estimated by the model-map Fourier Shell Correlations (FSC) curves using program *phenix.mtriage* (*34*). It is noteworthy that CryoFEM significantly augments the FSC resolution in comparison to the other methods. Intriguingly, irrespective of whether a soft mask (derived from the deposited PDB model) is used for calculating the FSC curves, CryoFEM yields identical results. This implies that CryoFEM can efficiently suppress background noise and does not require a soft mask for map enhancement. The box-whisker plots providing additional statistical information for all eight metrics reported here are available in Figs. S3 and S4. Fig. 2 and Fig. S5 are scatter plots for metrics in Table 1 and 2 across all 36 datasets. CryoFEM consistently enhances map quality across the tested samples.

**Figure 2:**
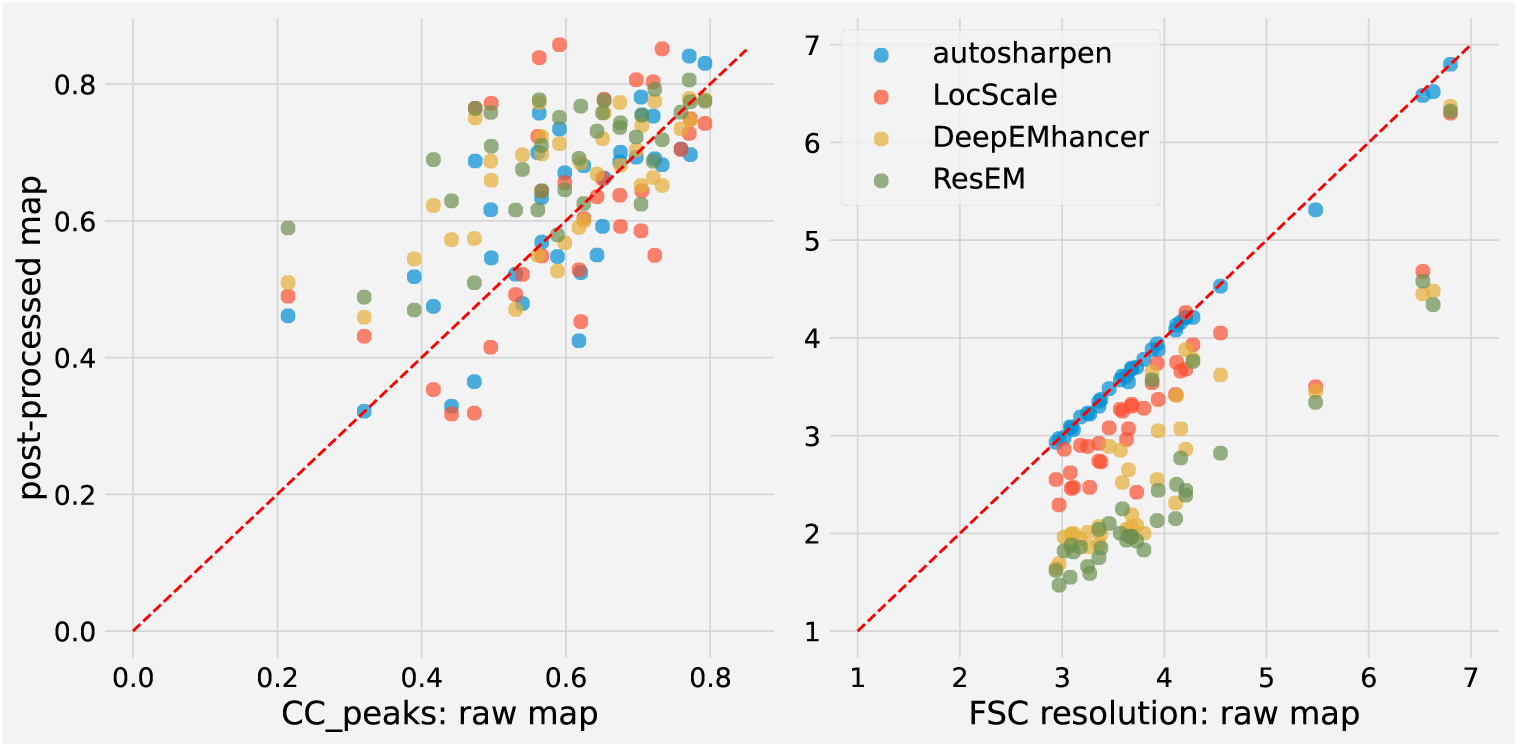
Distribution of CC_peak_ and FSC estimated resolutions. The scatter plots show the distribution of CC_peak_ values (left) and FSC resolution estimates at the 0.143 cutoff (right, unmasked 0.143) for the 36 test datasets. Each point represents the performance of a postprocessed map with respect to its raw map.

**Table 1:** Real space benchmark test. Median values of four model-map correlation coefficients in the real space on the test dataset (see Methods section for the definition of various correlation coefficients).

| Method | CC<sub>box</sub> | CC<sub>mask</sub> | CC<sub>volume</sub> | CC<sub>peak</sub> |
| --- | --- | --- | --- | --- |
| Raw | 0.738 | 0.728 | 0.737 | 0.608 |
| auto_sharpen (33) | 0.727 | **0.775** | **0.761** | 0.666 |
| LocScale (26) | 0.764 | 0.727 | 0.735 | 0.640 |
| DeepEMhancer (32) | 0.721 | 0.699 | 0.710 | 0.683 |
| CryoFEM | **0.808** | 0.731 | 0.739 | **0.714** |

**Table 2:** Fourier space benchmark test. Median resolutions were estimated by FSC curves using different cutoff and masking conditions for the test datasets. The masks were generated using the deposited PDB models.

| Method | Masked (0.143) | Unmasked (0.143) | Masked (0.5) | Unmasked (0.5) |
| --- | --- | --- | --- | --- |
| Raw | 3.27 | 3.67 | 3.69 | 4.42 |
| auto_sharpen (33) | 3.36 | 3.69 | 3.71 | 4.49 |
| LocScale (26) | 3.14 | 3.26 | 3.72 | 3.92 |
| DeepEMhancer (32) | 2.41 | 2.42 | 3.96 | 3.98 |
| CryoFEM | **2.02** | **2.02** | **3.40** | **3.40** |

### Map-Model building and refinement

We have established CryoFEM’s proficiency in revealing high-resolution features in cryo-EM maps. In this section, we explore whether this enhancement in map features can aid in the building and refinement of protein models. Our key assessment metrics are the C*_α_* score, GDTHA score, and seq-score (see Methods section for details). Each metric offers a measure of agreement between the predicted model and the deposited PDB model (Fig. 3a-c). In real-life situations, employing these metrics may not be feasible due to their dependence on the availability of a ground truth model. As a practical alternative, we compared correlations between the model and raw map (CC_volume_) in Fig. 3d. These evaluations provide a comprehensive view of the performance and impact of CryoFEM on protein model building and refinement. In Fig. 3a, we present the cumulative progression of the C*_α_* score for 17 selected datasets after a single cycle of our workflow shown in Fig. 1. Furthermore, Fig. 3b-d displays a comparison of the initial AlphaFold model with the refined model in terms of the GDT-HA score, seq-score, and model-map correlation. We selected datasets based on the following criteria: 1) data deposited after July 2020, when the training data of the AlphaFold was created; 2) remove data with a sequence identity of greater than 30%; 3) cryo-EM map resolutions falling within and distributed evenly across the range of resolutions in the training set; and 4) initial AlphaFold predicted models of low accuracy (samples with C*_α_* scores in the lower 50 percentile) to emphasize the value of the refinement process. Due to the constraints of these criteria, we also included one dataset, 6RTC, deposited in 2019 but with a low C*_α_* score. The 17 datasets chosen serve to demonstrate the effectiveness of our approach and highlight its limitations. Moreover, we selected the longest chain to model whenever possible for each PDB entry of a multi-protein complex. Consequently, the average number of protein residues is 1017. The refined models revealed a significant improvement over the AlphaFold models in most instances, with PDB entries 7MQS and 7NXD being notable exceptions. We attribute this failure to improve to a combination of low resolution (4.4 and 4.6 Å) and a suboptimal initial AlphaFold model (C*_α_* score of 0.4 and 0.25).

**Figure 3:**
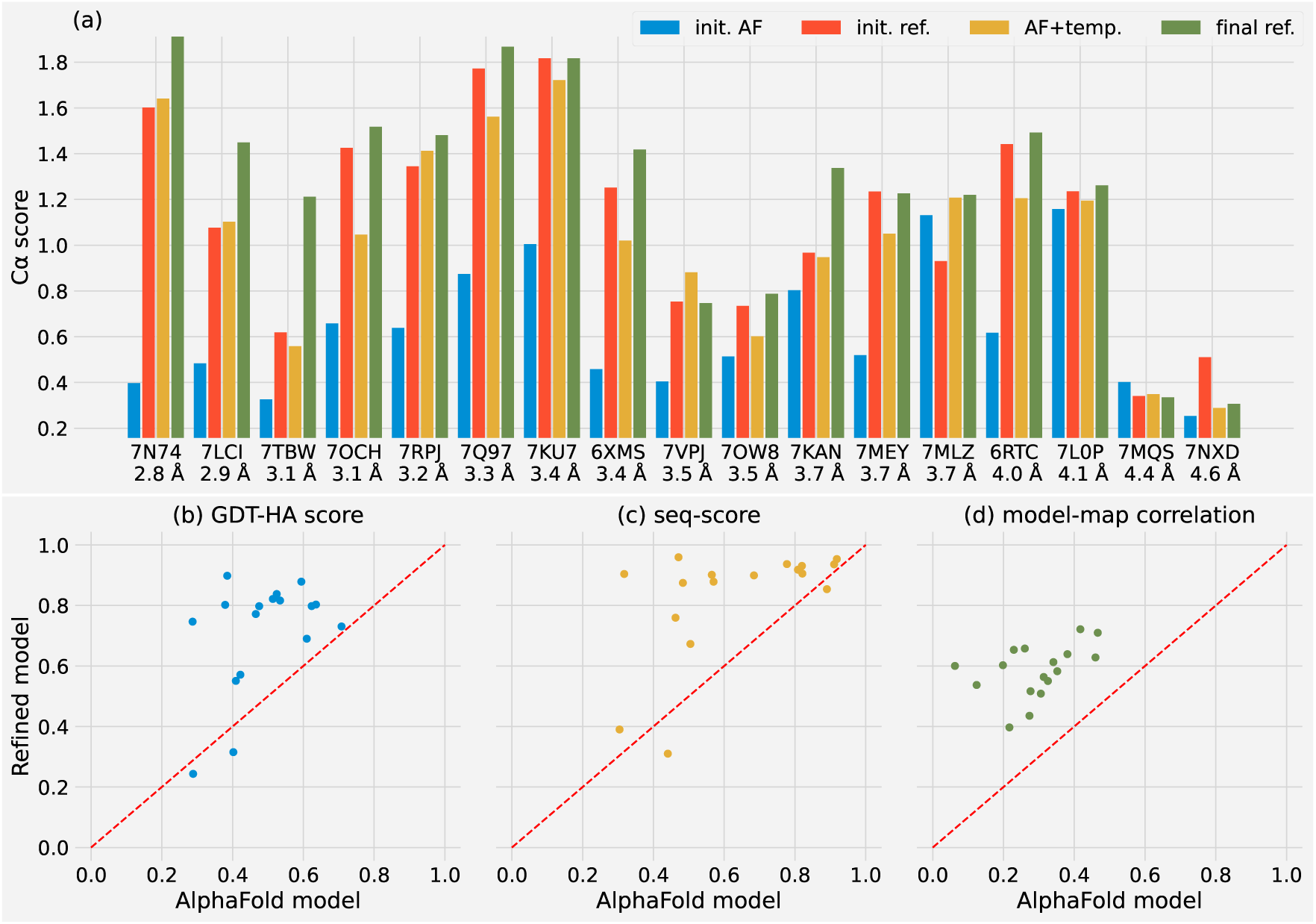
Model quality comparison between AlphaFold models and refined models. **(a)** Progression of C*_α_* scores per iteration. “init. AF” refers to the initial AlphaFold prediction; “init ref.” represents the first model-map refinement, with the refined model serving as a template in the “AF + temp” prediction. In “final ref.”, the predicted model undergoes refinement against the density map once more. **(b-d)** Comparison of the final refined model (y-axis) and the initial AlphaFold model (x-axis) in terms of **(b)** GDT-HA scores, **(c)** seq-scores, and **(d)** model-map correlations.

To underscore the value provided by our workflow, we conducted additional benchmark tests by substituting CryoFEM-enhanced maps with raw maps in the process. The outcomes are presented in Fig. 4. We noted that for PDB entries 6XMS and 7NXD, the second map-model rebuilding failed when utilizing raw maps. So, we reported the results from the preceding step (AlphaFold + template) instead for the two failed tests. In all 17 cases, our workflows leveraging CryoFEM-enhanced maps achieved higher C*_α_* scores compared to those using raw maps. On average, CryoFEM contributed to a 30.2%, 14.0%, 7.6%, and 17.4% increase in terms of C*_α_* score, GTS-HA score, seq-score, and model-map correlation, respectively. As can be seen in Fig. 3 and 4, CryoFEM-enhanced cryo-EM maps result in superior model quality.

**Figure 4:**
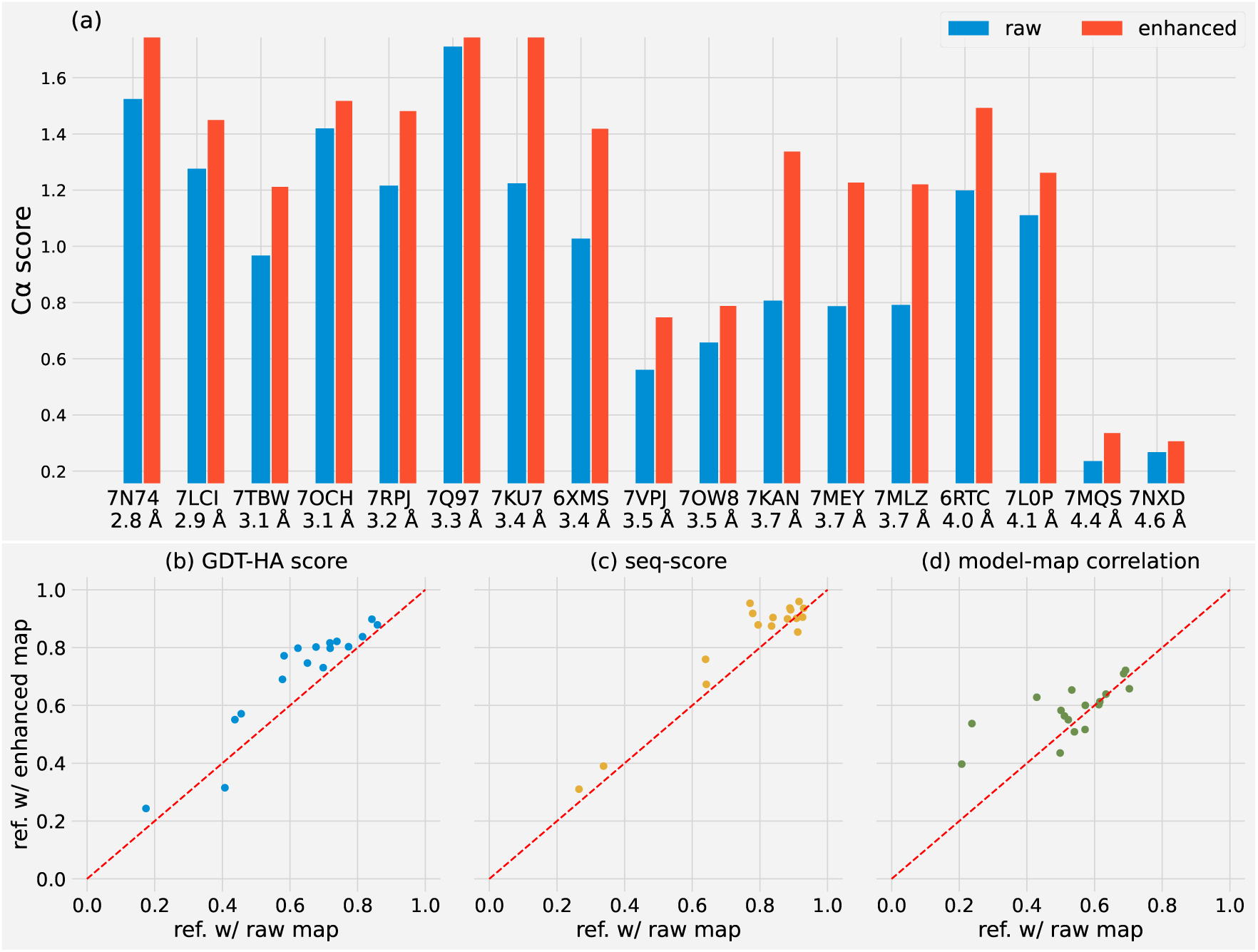
Benchmark comparison using raw maps. **(a)** C*_α_* scores depict the final refinement outcome after completing a full cycle. **(b-d)** Comparison of models refined with enhanced maps (y-axis) against those refined with raw maps (x-axis), focusing on **(b)** GDT-HA scores, **(c)** seqscores, and **(d)** model-map correlations.

To visualize the performance of our workflow, we present three samples, each with a different resolution (Fig. 5). In the first two cases (PDB entries 7TBW and 7OW8), *phenix.dock and build* managed to correct the poor AlphaFold models. Subsequent refinement using the CryoFEM-enhanced maps shows better results than those using the raw maps. PDB 7TBW is human ABCA1 cholesterol transporter bound with ATP (*35*). The density for the lid domain is not clear in the deposited cryo-EM map. After CryoFEM enhancement, the lid domain can be better visualized (Fig. 5a). PDB 7OW8 is a multidrug ABC transporter from Bacillus subtilis (*36*). The CryoFEM-enhanced map shows improved densities for loop regions (Fig. 5b). PDB 7L0P is an activated GPCR-G protein complex (*37*). In the deposited PDB, the AHD domain (*α*-helical domain) is missing due to poor density. Nevertheless, after CryoFEM enhancement, the density was recovered and the domain can be fitted in as a rigid body (Fig. 5c). The three examples demonstrate the utility of the CryoFEM to enhance cryo-EM maps for atomistic interpretation.

**Figure 5:**
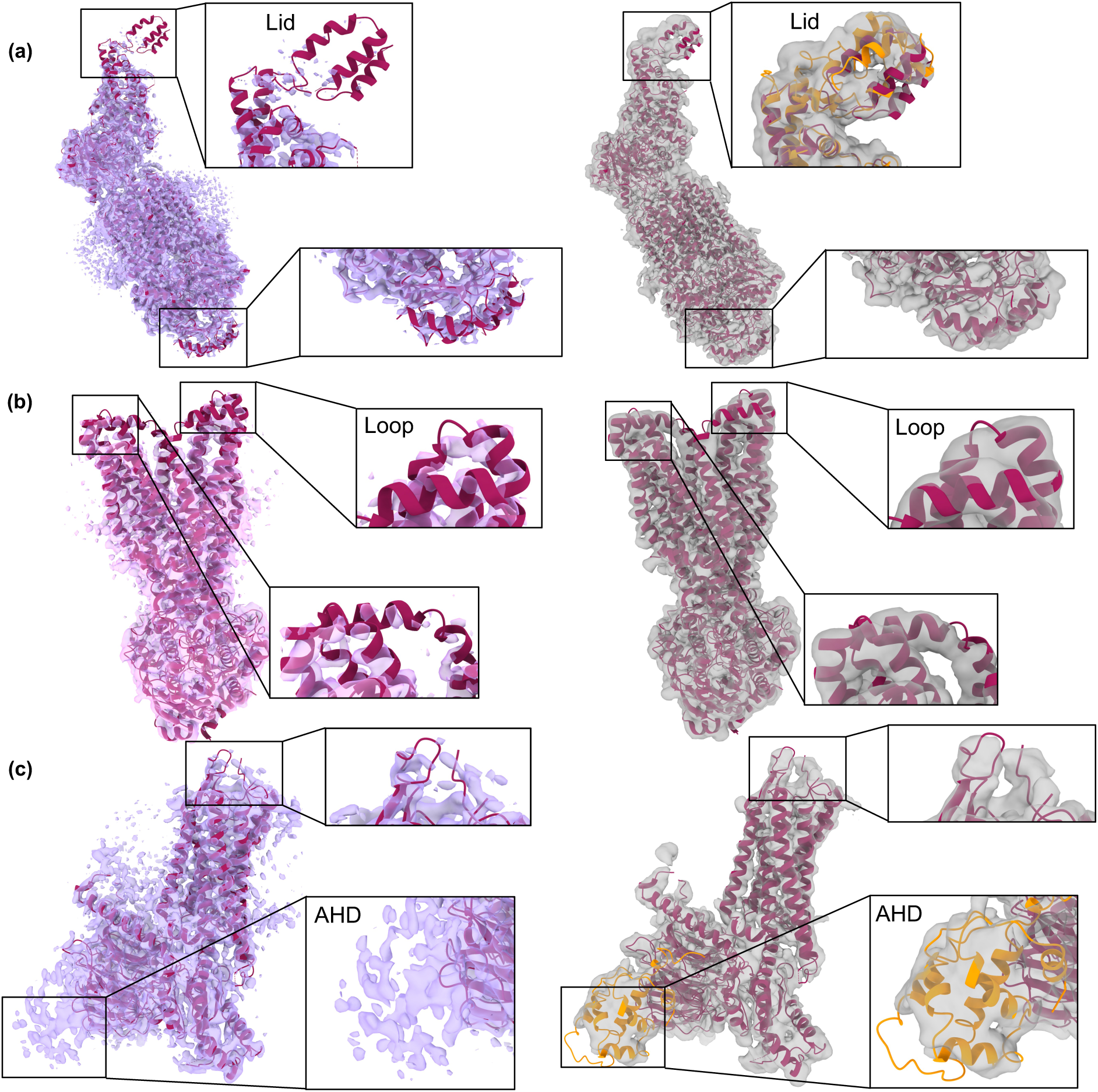
Comparative visualization of model support across different maps. Left column: deposited maps at author recommended contour level. Right column: CryoFEM-enhanced map. **(a)**: EMD-25800 (PDB entry 7TBW). **(b)** EMDB-13095 (PDB entry 7OW8) **(c)** EMD-23099 (PDB entry 7L0P). Maroon: deposited model. Orange: models from our automatic building pipeline.

## Discussion

In this study, we designed a map to model workflow by combining AlphaFold, deep learning image enhancement, and PHENIX map-model building tool to facilitate cryo-EM map interpretations. We tested the refinement progress for cryo-EM datasets with reported resolutions ranging from 2.8 to 4.6 Å (Fig. 3).

In most cases, our workflow improves the structure models significantly after one iteration. To achieve such improvement, we developed a deep learning model, CryoFEM, which enhances the atomic features of 3D cryo-EM maps, as evidenced by the improvement of real- and Fourier-space model-map correlations (Table 1,2). The CryoFEM-enhanced maps also benefit the PHENIX model building tool to refine AlphaFold predicted models (Fig. 4). Only for cryo-EM maps at low resolutions (*>* 4.4 Å) with very poor AlphaFold models (C*_α_* score lower than 0.4), our method failed to make an improvement to the structure model, which sets a current limit of applicability of our workflow.

The current workflow may be further optimized. After a full cycle, the resulting model can serve as the starting point for additional refinement by using *phenix.real space refine* and *Coot* to adjust the model-map fitness while maintaining good stereochemistry. Fig. 6 shows the realspace refinement results with *phenix.real space refine* for the same 17 PDB entries of Fig. 4. We can see that using CryoFEM-enhanced cryo-EM maps also helps with refinement and model building.

**Figure 6:**
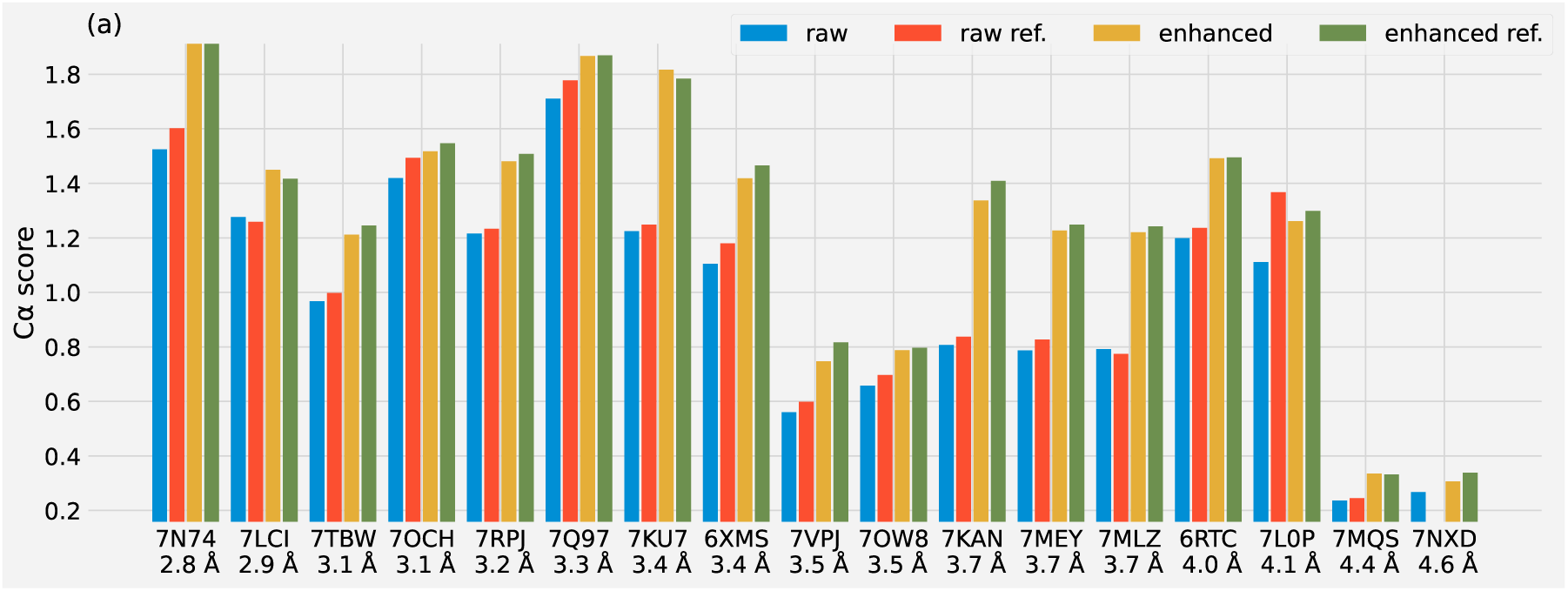
Model accuracy progress with *phenix.real space refine*. Here “raw” and “enhanced” refer to the models obtained after one full cycle of our workflow outlined in the Results session. “ref.” means refinement using *phenix.real space refine*. For 6XMS and 7NXD where the 2nd model-map rebuilding failed when using the raw map, we use the first rebuilt models for the refinement.

Through our experiments, we found that *phenix.dock and build* is sensitive to the map quality and the resolution limit at which it conducts the model building and refinement. Whenever using *phenix.dock and build* with CryoFEM-enhanced maps, we suggest trying to perform separate model building and refinement at the reported resolution and 2 Å, then select the model with a better model-map correction to proceed. When using the raw cryo-EM maps for model building and refinement, we set the resolution limit as the reported resolution.

Before our work, DeepEMhancer represented a popular deep-learning tool for post-processing cryo-EM density maps to recover high-resolution features. We managed to outperform DeepEMhancer (Tables 1, 2) while maintaining a similar model size (model parameters: 49.6 M (CryoFEM) vs 51.1 M (DeepEMhancer)). Notably, CryoFEM also significantly outpaces DeepEMhancer in terms of post-processing time: it took approximately 24 minutes for CryoFEM to process 36 maps in the benchmark test set, compared to the over 8 hours required by DeepEMhancer using identical hardware.

The efficacy of our CryoFEM model in enhancing map quality can be primarily attributed to two key factors. First, the model’s design is based on an augmented U-Net structure, which incorporates intra-block skip connections that facilitate hierarchical feature learning. This structure allows the model to capture both local and global contexts, which is demonstrated by the consistent FSC resolution results (Table 2), irrespective of the usage of the mask. Second, during training, we employed a composite loss function that combines L1 loss and a correlation loss (see Methods for details). This composite loss function ensures our model minimizes absolute discrepancies while preserving spatial relationships, thus enhancing the model’s robustness against the typical noise and outliers in cryo-EM maps.

There is understandable caution within the community regarding the use of deep learning post-processed maps for atomic model refinement, primarily due to the risks of over-fitting and hallucination. To address these concerns, we have taken deliberate steps to evaluate the model quality using raw maps and to demonstrate its robustness against noise contamination. Specifically, our model has been rigorously tested against both pure Gaussian noise (Fig. S3) and partial noise injection (Fig. S4). In the former scenario, the model effectively suppressed pure noise, while in the latter, it accurately identified and suppressed injected noise in a large contiguous block. These tests, along with extensive benchmarking, have shown significant improvements in real-space and Fourier space correlations (Fig. 2). By employing a carefully designed deep learning model and rigorous validation procedures, we have effectively mitigated the risks of over-fitting and hallucination. While acknowledging the importance of thorough validation, we believe that our approach represents a valuable advancement in cryo-EM map post-processing, enabling enhanced efficiency and accuracy in downstream model building and refinement tasks.

## Materials and Methods

### Data collection and preprocessing

Our training and testing datasets were sourced from the Electron Microscopy Data Bank (EMDB) (*38*). We set specific criteria for the selection process, focusing on entries with a reported resolution range of 2.5 to 5 Å and deposited between 01/01/2020 and 03/16/2023. We opted for entries providing valid half-maps, which allowed us to access the “unpolished” raw maps. To refine the selection further, we implemented pairwise sequence alignment using CD-HIT (*39*) to eliminate structures with sequence identity scores above 30%. A complete list of EMDB entry IDs associated with our study can be found in our GitHub repository (*40*).

We obtained the original raw map by averaging two half-maps. We then standardized the average map to a voxel volume of 1 Å^3^ using ChimeraX (*41*). The target map for training was derived from the corresponding PDB using the ChimeraX *molmap* command, with a resolution cutoff at 2 Å. We applied min-max normalization to all input and target maps to ensure standardization to a range of [0, 1]. Subsequently, we examined the Pearson correlations between the simulated and raw maps, removing entries with correlations lower than 0.15.

The final training set consisted of 1082 cryo-EM maps, with an additional 117 maps forming the validation set for monitoring training progress. We also reserved a test set of 36 cryoEM maps, including 20 entries previously used by DeepEMhancer. This test set was used exclusively for the examples presented in the Results section.

### Benchmark methods used in Table 1 & 2

Our benchmark test leveraged several existing tools, ensuring a robust and comprehensive evaluation of our method. As a representative of global-B factor sharpening methodologies, we employed *p*henix.auto sharpen (*33*). This method requires both the raw map and resolution parameters as inputs; for the latter, we used the resolution reported for each corresponding map to compute the sharpened output.

In addition to global-B factor sharpening, we also applied local B-factor sharpening via LocScale in its protein model-free mode (*26*), which can generate a pseudo-atomic model based on the input raw map and subsequently estimates the local B-factor. Using this method, we were able to produce corrected maps for 34 of 36 test samples.

Lastly, we used DeepEMhancer to test our workflow. DeepEMhancer necessitates two halfmaps as input. We opted to employ its default model weight and normalization mode (*32*). This mode has been designed to automatically estimate the noise level and normalize input maps, thus ensuring they have a zero mean and a standard deviation of 0.1.

### Map and model evaluation

#### Map quality

We used *phenix.map model cc* to evaluate real space correlations between the map and the corresponding deposited PDB model. In addition to the raw map, *phenix.map model cc* requires the resolution parameter, for which we used the reported resolution across all post-processed maps.

We used *phenix.map model cc* to perform analysis yielding four distinct correlation coefficients. CC_box_ represents the correlations across the entirety of the input map volume and the model-generated map. As such, this metric tends to be higher than the others in instances where large featureless regions exist. CC_mask_ calculates correlations using grid points selected by a molecular mask, as defined in Ref. (*42*). CC_volume_, on the other hand, employs a different mask that corresponds to the largest N grid points of the model map. For CC_peak_, the mask is similar to that used in the CC_volume_ computation, with the key distinction that it considers both the model map and the input map. The mask represents the union of the highest N points in both the model map and the input map.

To estimate map resolution using FSC curves, we deployed *phenix.mtriage*, which computes the model-map FSC curves. As highlighted in Ref. (*34*), the resolutions obtained from this approach align well with those derived from half-map correlations, when the cutoff is set to be 0.143. In instances of masked FSC calculation, *phenix.mtriage* applies a soft mask, derived from a binary mask used for CC_mask_, to circumvent Fourier artifacts.

#### Model quality

In this study, we primarily leverage the C*_α_* score, GDT-HA score, and seq-score to gauge the quality of our model against the deposited PDB model. To get C*_α_* and seq-score, we first employ *phenix.superpose and morph* to perform secondary structure matching (SSM), superimposing our output model with the deposited PDB model.

The C*_α_* score was calculated as the ratio of the proportion of “close” C*_α_* atoms to the RMSD (root mean square deviation) of these “close” C*_α_* atoms between the query and target PDB models. Here, two C*_α_* atoms are defined as “close” if their displacement is less than 2 Å after superposition. For example, if in a predicted model, 80% of the C*_α_* atoms are located within 2 Å of their corresponding atoms in the deposited model, and these atoms have an RMSD of 1.6 Å, the resulting C*_α_* score is 0.8/1.6 = 0.5. It’s important to note that the proportion is determined using the total number of C*_α_* atoms in the PDB model as the denominator when calculating the coverage.

Unlike the RMSD consideration, the seq-score is calculated as the product of the C*_α_* atom coverage (defined as the percentage of “close” C*_α_* atoms) and the percentage of sequences in the query model that match those in the target PDB model. Given that AlphaFold consistently predicts full sequence protein models, the sequence matching rate tends to be high, with an average sequence match of 91.1% for the 17 datasets in Fig. 3 from the initial AlphaFold predictions. Consequently, in our study, the seq-score primarily reflects improvements in terms of C*_α_* atom coverage. To generate the C*_α_* score and seq-score, we employed the tool *phenix.chain comparison* and set the *max dist = 2*.

The GDT-HA score represents the averaged C*_α_* atom coverage at 0.5, 1, 2, and 4 Å cutoffs. To obtain the GDT-HA score, we used TM align (*43*), which concurrently provides the TMscore and GDT-TS score. Results for these metrics are listed in Fig. 5. As TM-align conducts structure alignment prior to metric calculation, we provide the query model without utilizing *phenix.superpose and morph* in this case.

### Deep Learning Model

#### Model structure

CryoFEM ingests a raw cryo-EM map as input and outputs an enhanced map with matching dimensions. The network’s main components encompass encoding (down-sampling), bottleneck, and decoding (up-sampling) blocks. U-Net and its various derivatives have been shown to be proficient in image denoising tasks (*32, 44, 45*).

CryoFEM distinguishes itself from the classical U-Net through the integration of both inter- and intra-block residual connections. Inter-block connections facilitate hierarchical feature learning, capturing both local and global contexts that are critical for image enhancement. Simultaneously, intra-block connections address the vanishing gradients issue, thereby enabling the model to learn more features effectively (*46*). Furthermore, all convolutional layers in our architecture are bias-free, ensuring scale invariance. Our model employs learnable convolution layers, namely strided convolution and transposed convolution, for down-sampling and up-sampling processes, respectively (*44*). In each down-sampling block, the input data first traverses through the residual blocks before undergoing down-sampling via strided convolution. Similarly, during the up-sampling phase, the input data undergoes up-sampling via transposed convolution before passing through the residual blocks. The bottleneck block exclusively contains residual blocks. In our model, two residual blocks are assigned to each down-sampling, bottleneck, and up-sampling phase.

#### Training and validation of deep learning model

Given the varying dimensions of each cryo-EM map, we zero-padded and divided the map into smaller-sized chunks, for instance, 64^3^. This processed data was then reassembled to its original dimension for both the training and testing phases. To mitigate boundary artifacts, we employed a strategy as outlined in Ref. (*14*), utilizing only the central part (e.g., 50^3^) of the output without overlap during the reassembly stage.

Our initial experimentation involved a chunk size of 64; however, we observed that a size of 128 resulted in lower losses. Owing to the larger memory demand of 128^3^ chunks, which only permitted a smaller batch size of 2, thus slowing training, we opted for a two-stage training process. The model was first warmed up using the smaller chunk size of 64 for the initial four epochs before transitioning to the larger 128^3^size, with a central part of 100^3^utilized in the reassembly process.

Aiming for enhanced model generalizability, we augmented the input maps with random Gaussian noise, random anisotropy, and random blur using TorchIO (*47*) for the training data. Our loss function combines L-1 loss and real-space correlation. Specifically, the total loss amounts to *L*_1_(*y*_pred_*, y*_target_) + *w*(1 − corr(*y*_pred_*, y*_target_)), with corr(*y*_pred_*, y*_target_) denoting the Pearson correlation. During the warm-up phase, the combination weight, *w*, was set to 0.5 before being reduced to 0.1 in the later fine-tuning stage.

For optimization, we used AdamW (*48*) in both warm-up and fine-tune stages, with a batch size of 16 (16 ×64^3^ cubes) and 2 (2 ×128^3^ cubes), a weight decay of 1 × 10^−5^, and an initial learning rate of 8×10^−5^, which was decreased to 4×10^−5^ in the fine-tuning stage. We concluded the total training after 8 epochs when the validation loss began to rise, and we selected the model with the lowest validation loss at the 7th epoch. A detailed view of the L1 loss and correlations for each map throughout the entire training process is provided in Fig. S4.

Besides standard training parameters, the most influential hyperparameter was found to be the number of residual blocks in each stage of the model, namely, up-sampling, bottleneck, and down-sampling, (see Fig. S1 and Table S1 for a detailed depiction of the model structure). After experimenting with 1, 2, and 3 residual blocks, we determined that 2 blocks were optimal for the available training data. CryoFEM was implemented in PyTorch v2.0 (*49*), with all training and testing carried out on a single Nvidia V100 GPU. Warm-up and fine-tune required approximately 26 and 40 hours per epoch, respectively.

### AlphaFold prediction

In order to ensure adaptability and expandability within our workflow, we employ OpenFold (*50*), a faithful open-source implementation of AlphaFold, for protein structure predictions. Initial predictions are conducted using five distinct AlphaFold models (as detailed in Supplementary Table 5 of Ref. (*17*)), with the model yielding the highest pLDDT score selected as the initial model. After obtaining the template from the first Phenix model building, two subsequent runs are conducted – one with and one without multiple sequence alignment (MSA). Each run incorporates two AlphaFold models trained with the template, with the model delivering the highest pLDDT score being chosen from each run. Thereafter, the model (either with or without MSA) that yields the highest C*_α_* score is selected for the next iteration of Phenix model building. This iterative approach ensures a continual refinement of models leading to the best possible prediction.

### Model building and refinement

As delineated in Ref. (*23*), *phenix.dock and build* first discards the residues in the AlphaFold model with pLDDT values below 70. Subsequently, the model is segmented into multiple domains, each is individually docked into the density map. In all cases except for PDB entry 7TBW studied in Fig. 3 and 4, we employed the default parameters which split the model into a maximum of 3 domains. However, 7TBW, which is the largest model in our study (2270 residues) exhibits a considerably low initial AlphaFold model quality (C*_α_* score 0.33). We discerned that model rebuilding could benefit from increasing the maximum domains to 8 when using either the CryoFEM enhanced map or the raw map.

## Acknowledgments

We would like to thank Sean McSweeney and John Shanklin for the helpful discussion. This work was supported in part by Brookhaven National Laboratory LDRD 22-008. Q. L. was supported by the U.S. Department of Energy (DOE), Office of Biological and Environmental Research as part of the Quantitative Plant Science Initiative at BNL.

## Author contributions

X.D. and Q.L. designed the study and experiments. X.D. and L.W. performed the experiments. X.D., L.W., S.Y., and Q.L. analyzed the data. X.D. and Q.L. wrote the manuscript with help from other coauthors.

## Data and materials availability

All data used in this study are publicly available from https://www.rcsb.org (for PDB structures) and https://www.emdataresource.org for cryo-EM maps. CryoFEM is developed as an open-source Python package, freely available at https://github.com/Structurebiology-BNL/CryoFEM. This repository includes all codes necessary for training and testing. For enhanced accessibility and user-friendly interaction, we have also implemented an easy-to-use Google Colab notebook, which serves as an inference server. It can be accessed at https://colab.research.google.com/github/Structurebiology-BNL/CryoFEM/blob/main/Colab_CryoFEM.ipynb.

