## Supplementary Materials for "Integrating AlphaFold and deep learning for atomistic interpretation of cryo-EM maps"

### Model structure

Fig. S1 shows a schematic architecture of CryoFEM. For simplicity, we only show one residual block (ResBlock) for each downsampling, bottleneck and upsampling block. The model used in the main text employed 2 ResBlocks in each phase. Table S1 includes a detailed account

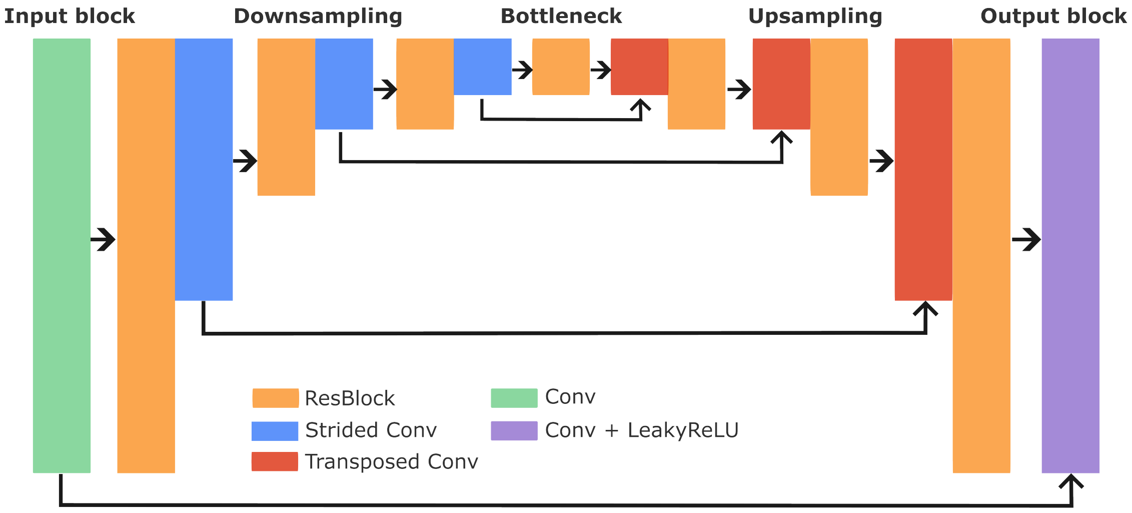


Figure S1: Schematic of CryoFEM model architecture. Black cornered arrows denote inter-block skip connections.

of the model components and the shapes of input and output per each block.

### Model training progress

Figure S2 presents the evolution of the $L_1$ loss and Pearson correlation coefficients per each map during the training and validation phases of our model over 8 epochs. The model checkpoint from the 7th epoch, which exhibits the lowest total loss $(L_1 + 0.1 * (1 - \text{pcc}))$, is chosen as the final model for the analysis in the main text.

| Module | Component | Input Shape | Output shape |
|---|---|---|---|
| Input block | Conv3D | (1, 128, 128, 128) | (64, 128, 128, 128) |
| Down-sample block 1 | 2 ResBlocks + Conv3D(stride=2) | (64, 128, 128, 128) | (128, 64, 64, 64) |
| Down-sample block 2 | 2 ResBlocks + Conv3D(stride=2) | (128, 64, 64, 64) | (256, 32, 32, 32) |
| Down-sample block 3 | 2 ResBlocks + Conv3D(stride=2) | (256, 32, 32, 32) | (512, 16, 16, 16) |
| Bottleneck block | 2 ResBlocks | (512, 16, 16, 16) | (512, 16, 16, 16) |
| Up-sample block 1 | ConvTranspose3D + 2 ResBlocks | (512, 16, 16, 16) | (256, 32, 32, 32) |
| Up-sample block 2 | ConvTranspose3D + 2 ResBlocks | (256, 32, 32, 32) | (128, 64, 64, 64) |
| Up-sample block 3 | ConvTranspose3D + 2 ResBlocks | (128, 64, 64, 64) | (64, 128, 128, 128) |
| Output block | Conv3D + LeakyReLU | (64, 128, 128, 128) | (1, 128, 128, 128) |

Table S1: Detailed model structure with the shape information. For the input $x$, the ResBlock will yield $x + \text{Conv3D}(\text{ReLU}(\text{Conv3D}(x)))$. Here for simplicity we have omitted the batch dimension.

### Robustness against corrupted input

In this section, we demonstrate the robustness of CryoFEM in handling noise-contaminated input maps. We specifically focus on two scenarios to validate the model's ability to suppress noise without introducing artifacts in the prediction. We first investigate a case where the input data consists entirely of random Gaussian noise with zero mean and a standard deviation of 1. As depicted in Fig. S3, our model successfully identifies and removes almost all the noise, outputting a map close to zero everywhere. The output data range is a minimal -5e-4 to 4e-3, in contrast to the input data range of -5.24 to 5.32, illustrating the model's effectiveness in handling pure noise.

Next, we present a more challenging scenario where only a partial portion of the input map (EMDB-13095) is replaced with Gaussian noise. Specifically, we randomly replaced a contiguous $40 \times 40 \times 40$ block of the input map with Gaussian noise of zero mean, and the standard deviation is set to 5 times that of the original map's standard deviation. Fig. S4 reveals that the model accurately identifies the corrupted portion and suppresses the injected noise.

These experiments underscore the robustness of our model in handling cases where the input map is deliberately injected with noises. The ability to correctly identify and suppress both pure and partial noise without introducing artifacts ensures that our model is a reliable tool for enhancing the quality of density maps in cryo-EM experiments.

### Detailed results of benchmark test

#### Model map correlation and FSC resolution

Fig. S5 and S6 shows the box-whisker plots for the real-space correlation coefficients and the FSC estimated resolution reported in the Table 1 and 2 of the main text. Fig. S7 shows the scatter plots of the three model-map real space correlations (upper panel) and three FSC resolutions (lower panel) using different post-processing methods for all 36 samples tested in

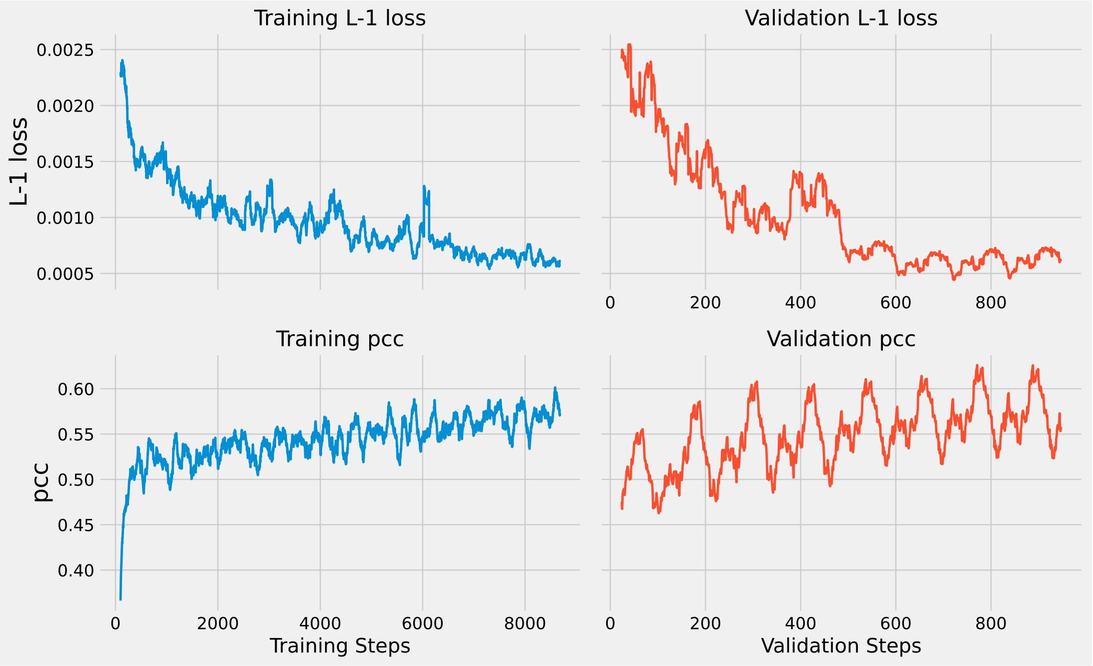


Figure S2: Training progress over 8 epochs. To smooth out the plots, we employ a moving average over every 100 maps for the training plots. Similarly, a 25-step moving average is applied to the validation plots.

the main text.

### FSC curves

Fig. S8 show Fourier Shell Correlation (FSC) curves for 8 samples across an array of map enhancement methods. Selected to represent a range of reported resolutions (2.8 Å to 4.4 Å), the samples undergo enhancement via methods including global B-factor sharpening (*phenix.auto_ sharpen*) and local-B factor sharpening (LocScale), DeepEMhancer, and CryoFEM. The correlations were computed using phenix.mtriage between the map and the corresponding PDB model. Their FSCs are juxtaposed with those of the raw map, demonstrating CryoFEM's effectiveness across different resolution levels.

### TM-score and GDT-TS score

Fig. S9 demonstrates the enhanced quality of models refined with CryoFEM enhanced maps, as evidenced by higher GDT-TS and TM-scoresscores compared to the original AlphaFold model and the model refined using the raw map. The models refined using CryoFEM enhanced maps yields an average improvement of 24.0% and 7.2% in GDT-TS and TM-scores, respectively, when compared with the original AlphaFold model. These improvements are still notable at 7.1% and 4.8% for GDT-TS and TM-scores, respectively, when compared with models refined with raw maps. Despite the modest improvements in TM-score, due to the already high TM-scores of the original AlphaFold models, the consistent enhancements across both metrics underscore the effectiveness of the CryoFEM refinement process.

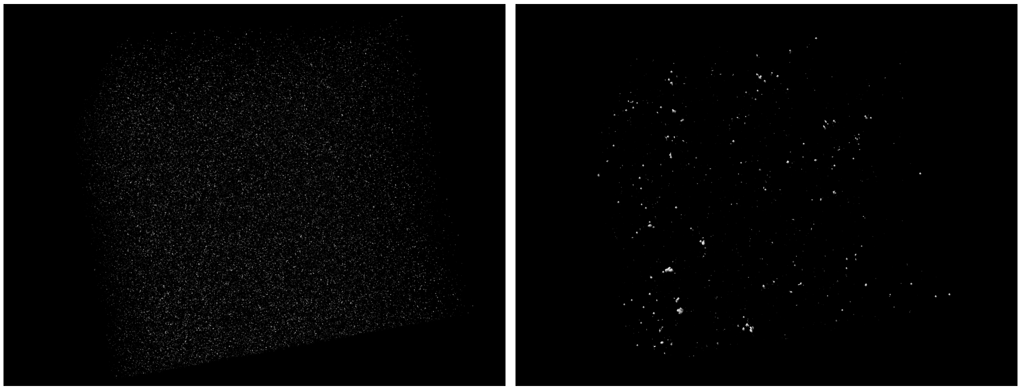

Figure S3: Left: The input map, consisting entirely of random Gaussian noise with zero mean and a standard deviation of 1. Right: The output map, where the model has successfully suppressed nearly all the input noises, resulting in values close to zero everywhere.

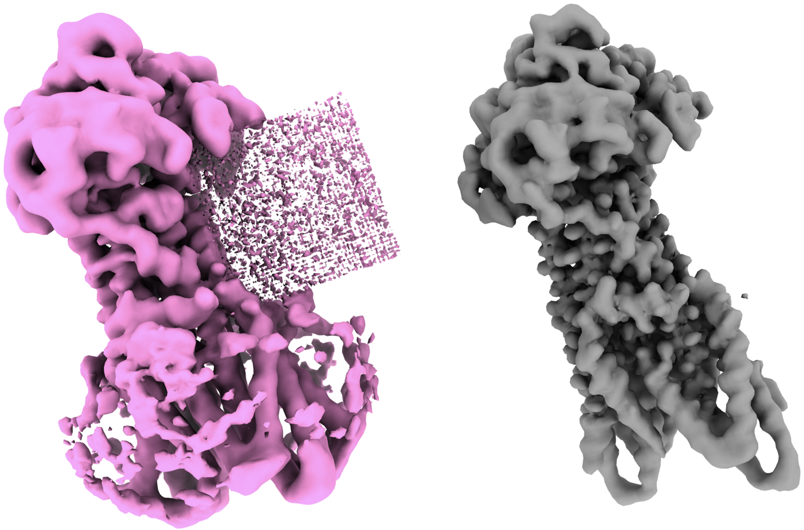

Figure S4: Left: The corrupted input map (EMDB-13095) with a $40 \times 40 \times 40$ contiguous block replaced by Gaussian noise, visible on the right side. Right: The predicted map, where the model has accurately identified and completely suppressed the injected noise.

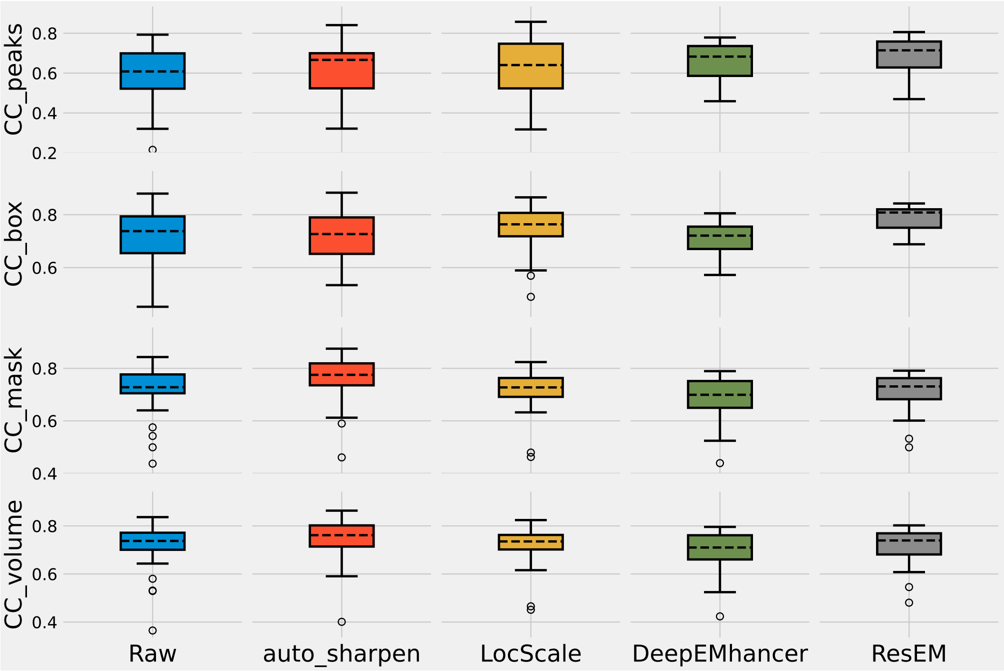


Figure S5: Real-space model-map correlations for different map processing methods on the 36 test maps. The box itself represents the interquartile range (IQR), from the first quartile (Q1) to the third quartile (Q3), with the black dashed line inside the box marking the median value. The whiskers extend from the box to cover the range within 1.5 times the IQR from the first and third quartiles. Hollow circles represent outliers defined as values that fall outside of this range.

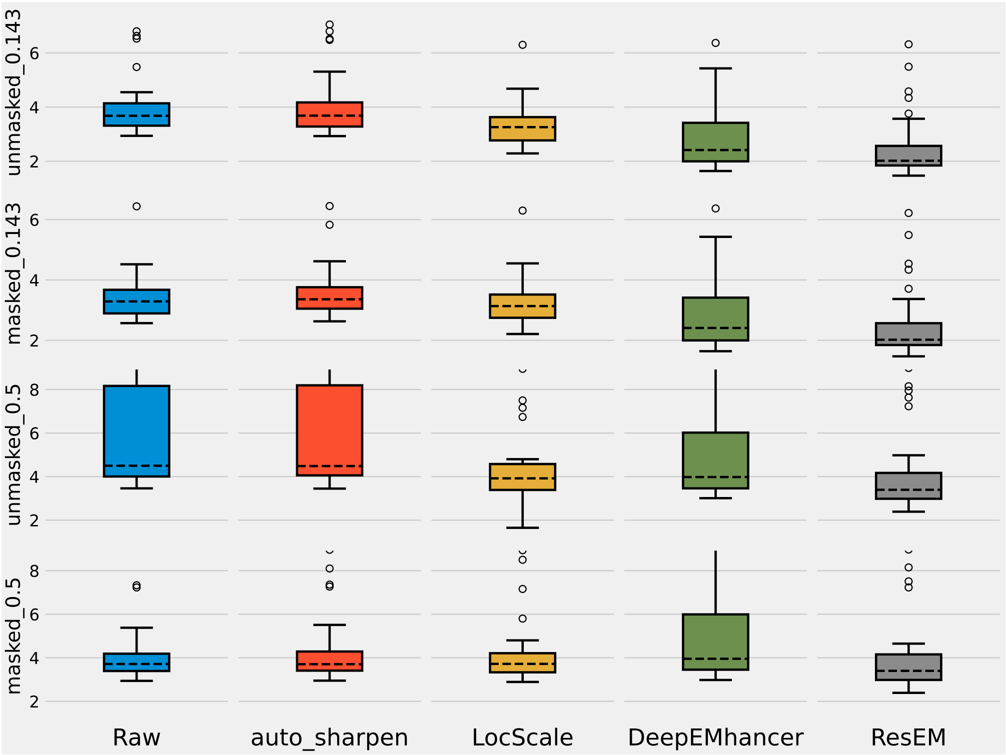


Figure S6: Resolutions estimated by model-map FSC curves for different methods on the 36 test maps. The box itself represents the interquartile range (IQR), from the first quartile (Q1) to the third quartile (Q3), with the black dashed line inside the box marking the median value. The whiskers extend from the box to cover the range within 1.5 times the IQR from the first and third quartiles. Outliers, defined as values that fall outside of this range, are represented by hollow circles.

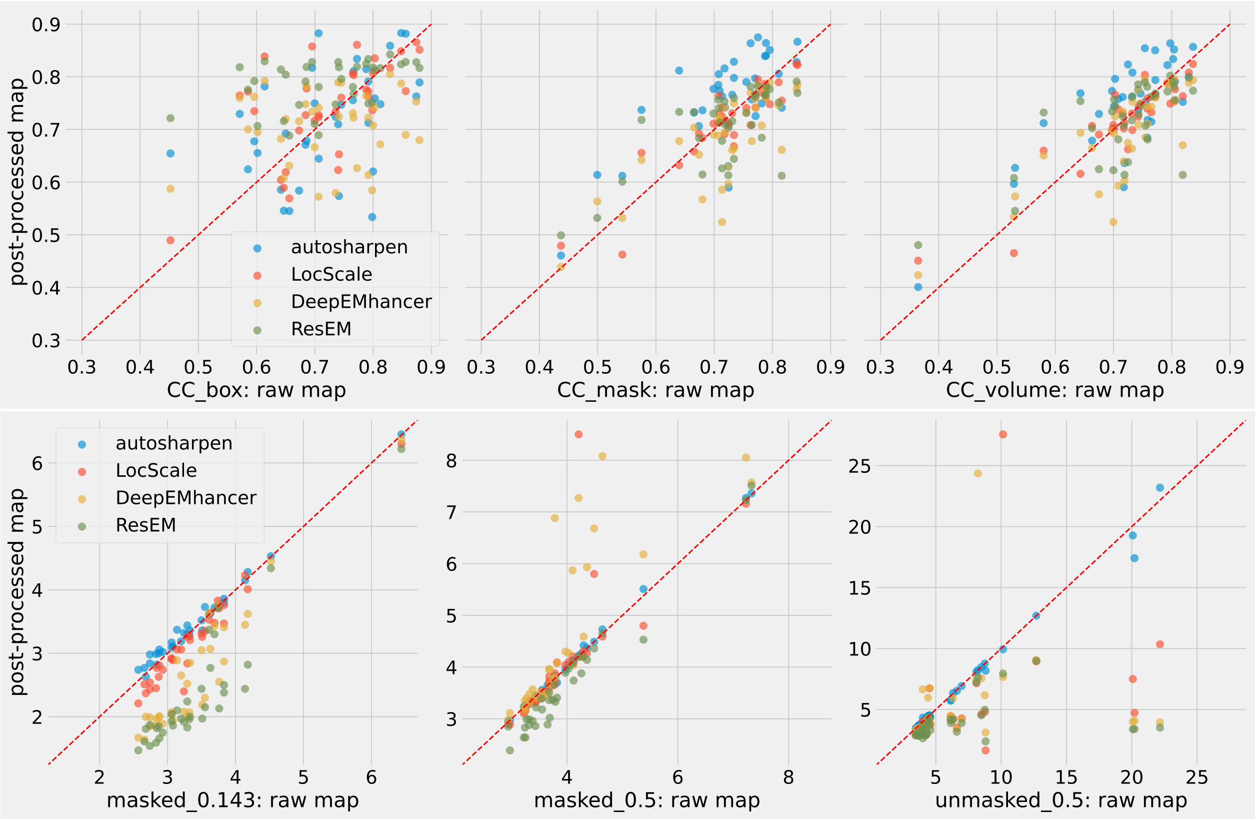


Figure S7: Upper panel: Real space model-map correlations (CC_box, CC_mask and CC_volume). Lower panel: Resolution estimated using model-map FSC curves calculated at different cutoffs (0.143 and 0.5) and masking (no-mask or soft-mask generated by the model).

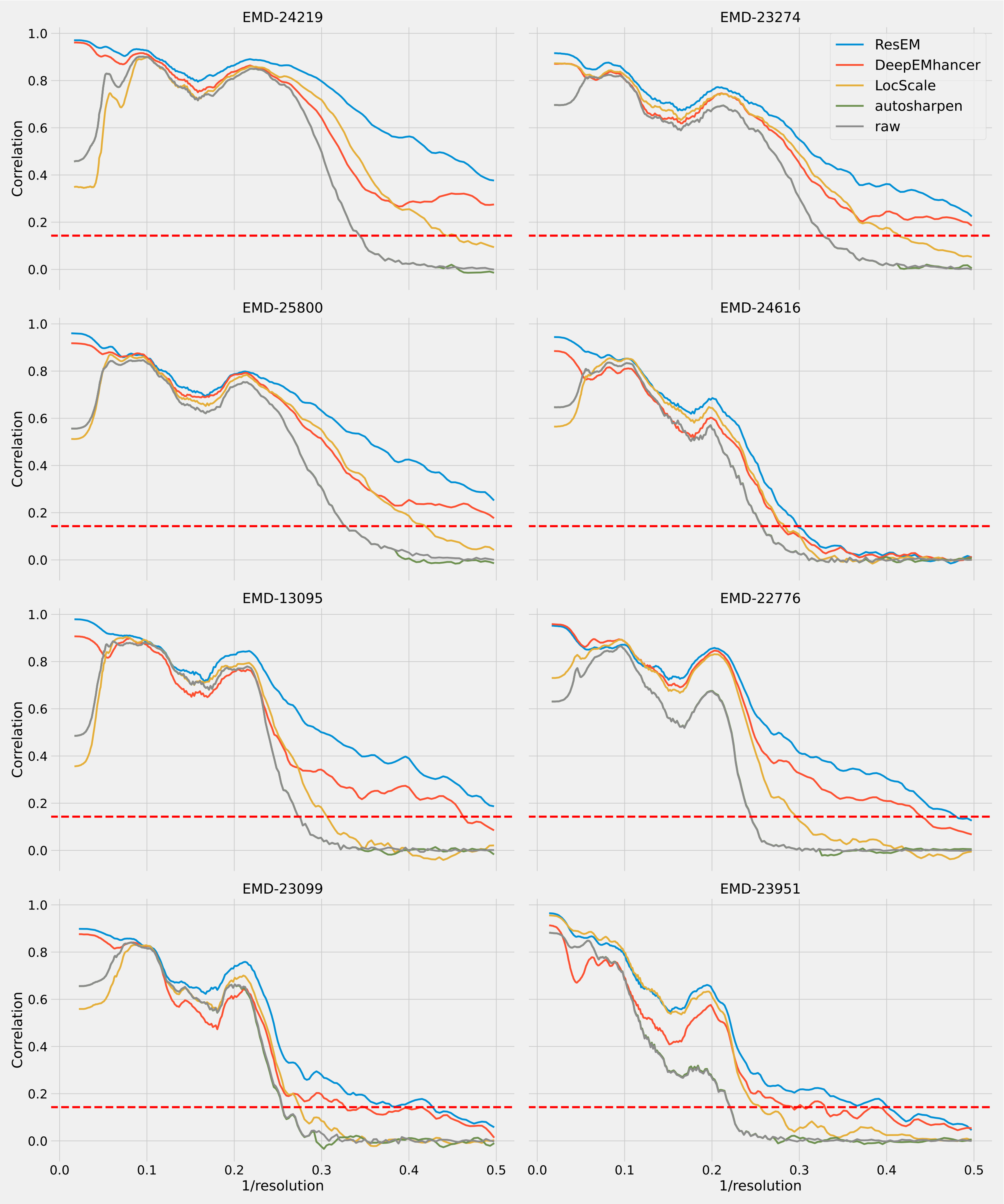


Figure S8: FSC curves (unmasked) for 8 samples with reported resolutions range from 2.8 Å to 4.4 Å. A reference line (red dashed) is drawn at a correlation of 0.143.

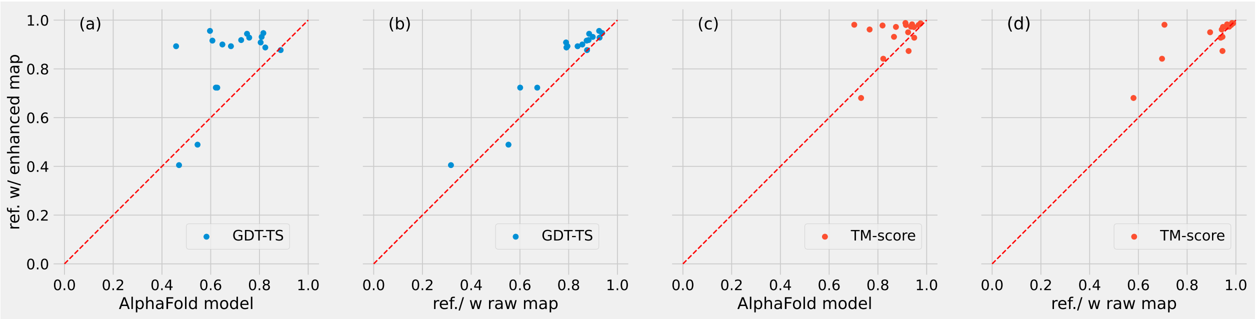


Figure S9: Comparative evaluation of model quality using GDT-TS score (blue) and TM-score (red). The y-axis shows the results obtained from models refined with CryoFEM enhanced map compared with the original AlphaFold model (a, c) and the model refined using raw maps (b, d).
